# CuiT is a Cu importer required for metal homeostasis in *Salmonella enterica*

**DOI:** 10.64898/2026.01.22.701185

**Authors:** Zhenzhen Zhao, Karla F. Díaz Rodríguez, Andrea A. E. Méndez, Lisandro M. Sommer, Pedro Mendes, Fernando C. Soncini, Susana K. Checa, Teresita Padilla-Benavides, José M. Argüello

**Affiliations:** Department of Chemistry and Biochemistry, Worcester Polytechnic Institute, 100 Institute Road, Worcester, MA 01609, USA; Instituto de Biología Molecular y Celular de Rosario, Facultad de Ciencias Bioquímicas y Farmacéuticas, Universidad Nacional de Rosario, Consejo Nacional de Investigaciones Científicas y Técnicas, Rosario, Argentina; Center for Cell Analysis and Modeling, University of Connecticut School of Medicine, Farmington, CT, 06030, USA; Department of Molecular Biology and Biochemistry, Wesleyan University, Middletown CT, 06459, USA

**Keywords:** Cu homeostasis, *Salmonella enterica*, CuiT, Cu-importer, Cu stress

## Abstract

Copper (Cu) is an essential micronutrient that serves as a cofactor for redox enzymes but becomes toxic when unregulated. In bacteria, while Cu efflux systems are well characterized, mechanisms of Cu import remain poorly understood. Here, we characterize the major facilitator superfamily transporter CuiT (STM1486) as a key Cu importer in *Salmonella enterica*. Comparative genomics revealed that *cuiT* is evolutionarily conserved across Enterobacteriaceae, and structural modeling predicts a 12-transmembrane-helix architecture with conserved His, Met, and Cys residues suitable for Cu coordination. Functional analyses demonstrated that deletion of *cuiT* reduces intracellular Cu accumulation, slows Cu uptake kinetics, and diminishes expression of Cu-responsive genes, including *copA*, *cueP*, *cueO*, and *golB*. Conversely, overexpression of CuiT increases intracellular Cu but sensitizes cells to Cu stress, highlighting the need for tight regulation. Kinetic modeling indicates that CuiT mediates rapid Cu import, supporting larger intracellular Cu pools compared to *Pseudomonas* influx transporters. These findings position CuiT as a central component of the *Salmonella* Cu homeostasis network, linking Cu import to transcriptional regulation, redox balance, and stress adaptation. Our work provides mechanistic insights into bacterial Cu acquisition and suggests CuiT and associated pathways as potential targets for antimicrobial strategies.

**Significance:** Copper (Cu) is essential for bacterial redox enzymes but toxic when dysregulated. While Cu efflux pathways are well studied, mechanisms of Cu import are poorly understood. We identify CuiT, a conserved major facilitator superfamily transporter, as a key Cu importer in *Salmonella enterica*. CuiT controls intracellular Cu levels, influences Cu-responsive gene expression, and maintains redox balance and stress adaptation. Disruption or overexpression of CuiT perturbs Cu homeostasis, highlighting its regulatory importance. These findings reveal a critical bacterial Cu acquisition pathway and suggest CuiT and its network as potential antimicrobial targets, advancing understanding of metal homeostasis in pathogens.

## INTRODUCTION

Copper (Cu) is an essential micronutrient that serves as a cofactor for electron transport proteins and redox enzymes, such as plastocyanin and cytochrome *c* oxidases (1–3). To ensure proper incorporation into cuproproteins, organisms have evolved diverse Cu^+^ distribution mechanisms, including transporters for translocation across lipid bilayers, soluble cuprochaperones for intracellular distribution, and Cu^+^ sensing transcription regulators that control homeostasis-related gene expression (1–6). These systems are crucial not only for delivering Cu^+^ to target proteins but also for preventing toxicity associated with free Cu^+/2+^ ions, which can generate reactive oxygen and nitrogen species (ROS/RNS), oxidize thiol groups, and displace essential metals by binding to non-cognate sites (7, 8).

Transmembrane Cu^+^-transporters play a central role in maintaining cellular Cu levels (1, 4, 9). Cu^+^-ATPases, found across all domains of life, drive cytoplasmic efflux of the ion into extracellular or organellar compartments (9–11). Many bacteria also employ the tripartite CusABCF system for Cu^+^ efflux (12, 13). Deletion of bacterial Cu^+^ efflux systems result in easily noticeable metal sensitivity, slow growth and cell death (1, 4). However, identifying Cu^+^ influx transporters has been more challenging, as their removal does not yield clear phenotypic changes. Nonetheless, multiple Cu^+/2+^ import pathways have been proposed in both Gram-negative and Gram-positive bacteria (14). For instance, *Rhodobacter capsulatus* CcoA, a transporter member of the major facilitator superfamily (MFS), functions as a Cu^+^ importer required for cytochrome *c* oxidase activity, with conserved methionine (Met) and histidine (His) residues facilitating intracellular Cu^+^ allocation (15). In *Bacillus subtilis*, the YcnJ transporter, along with the putative Cu^+^-binding protein YcnI and the transcriptional repressor YcnK, are implicated in Cu^+^ uptake (16–18). Other proposed Cu^+^ importers include *Enterococcus hirae* CtaA and *B. subtilis* ZosA, though further validation is needed (19, 20). Additionally, porins contribute to Cu^+/2+^ influx, as observed in *Mycobacterium smegmatis*, where MspA and MspC mutants exhibit impaired Cu uptake (21–23). In *Pseudomonas stutzeri*, the TonB-dependent NosA mediates Cu uptake for nitrous oxide reductase assembly (24, 25), while its *P. aeruginosa* OprC, binds Cu^2+^ and its expression is repressed under Cu^2+^ stress (26, 27). In *Escherichia coli*, the outer membrane protein ComC was identified as a general stress response protein, although mutation of *comC* results in lower Cu^2+^ tolerance and reduced Cu^+/2+^ import (28, 29).

We have characterized the Cu^+^ homeostasis mechanisms in the opportunistic pathogen *P. aeruginosa* (30–34). This organism possesses well-defined efflux transporters (CopA1, CopA2, and CusABCF), influx transporters (OprC and CcoA), and Cu^+^-sensing transcriptional regulators (CopRS and CueR). Transcriptome analyses following Cu^2+^ stress revealed upregulation of Cu^+^ efflux transporters and downregulation of OprC (31). Interestingly, expression of CcoA the Cu^+^ importer involved in cytochrome C oxidase metalation remained unchanged, while the novel MFS transporter CuiT (PA5030) was repressed, hinting at a possible role in metal influx (31).

In this study, we investigate CuiT (STM1486) in *Salmonella enterica*, a major foodborne pathogen responsible for over 27 million infections and 200,000 deaths annually (35). Transition metals, particularly Cu^2+^, influence host-pathogen interactions, playing a central role in the oxidative burst as part of the innate immune response (6, 36). The Cu^+^ homeostasis network in *Salmonella* includes the transcriptional regulator CueR controlling the expression of the Cu^+^-ATPase CopA, a periplasmic chaperone CueP, and the multicopper oxidase CueO (3). Additionally, GolS, a CueR-like sensor, regulates GolB the sole apparent cytoplasmic Cu^+^ chaperone, and the Cu^+^-ATPase GolT (3). Unlike many other enteric species, the *Salmonella* core genome lacks the genes coding for the CusCFBA efflux system and seems to rely on CueP to compensate for the absence of the three partite efflux system (37, 38). CueP expression requires the joint activation of CueR and the envelope stress-induced two-component CpxRA system (39). Also, CpxRA triggers the expression of the Cu-activated ScsABCD system, comprising thiol-oxidoreductases and membrane-linked proteins, linking Cu^+^ homeostasis to redox stress resistance (6, 40). Despite these insights, Cu^+/2+^ import mechanisms remain poorly understood (3, 5).

To characterize *Salmonella* CuiT, we examined bacterial Cu^2+^ tolerance, gene expression, and Cu uptake in wild-type, CuiT-overexpressing, and *cuiT* deletion mutant *Salmonella* strains. CuiT expression was reduced under Cu stress and CuiT overexpression reduced tolerance to this ion, while *cuiT* deletion impaired Cu^+/2+^ uptake. Sequence analysis combined with structural modeling with Alpha Fold revealed conserved Met, His, and cysteine (Cys) residues in CuiT transmembrane fragments, congruent with Cu^+^ transport function. These findings suggest CuiT facilitates periplasm-to-cytoplasm Cu^+^ movement, broadening our understanding of bacterial Cu import systems.

## RESULTS

### Evolutionary conservation and structural modeling insights into CuiT-mediated Cu-transport

Conservation of a gene and its genomic environment provide information on its likely functional importance. To this end, we performed a comparative genomic analysis of chromosomal regions harboring *cuiT* homologs (**Figure 1A**). The analysis illustrates the synteny of *cuiT* and its neighboring genes across a diverse set of bacterial taxa (41). The analysis revealed that *cuiT* is flanked by a highly conserved gene coding for a LysR family regulator across different genera, with a strong syntenic conservation in *S. enterica* and closely related *Enterobacteriaceae* species, such as *E. coli* and *Shigella flexneri*. Interestingly, *cuiT* and the *lysR*-like homolog are also present in more distantly related bacteria, including *P. aeruginosa*, *B. subtilis*, and *Burkholderia cenocepacia*. This widespread distribution across phylogenetically diverse bacterial species suggests that the role of CuiT in Cu transport may be evolutionarily conserved beyond *Salmonella*. Conversely, in more distantly related bacteria such as *Mycobacterium avium* and *Corynebacterium glutamicum*, the genomic context surrounding *cuiT* appears more divergent, possibly reflecting species-specific adaptations in Cu regulation. These findings support the hypothesis that CuiT is a key component of Cu homeostasis pathways across multiple bacterial lineages, with potential functional conservation in Cu uptake and stress response mechanisms.

**Figure 1.**
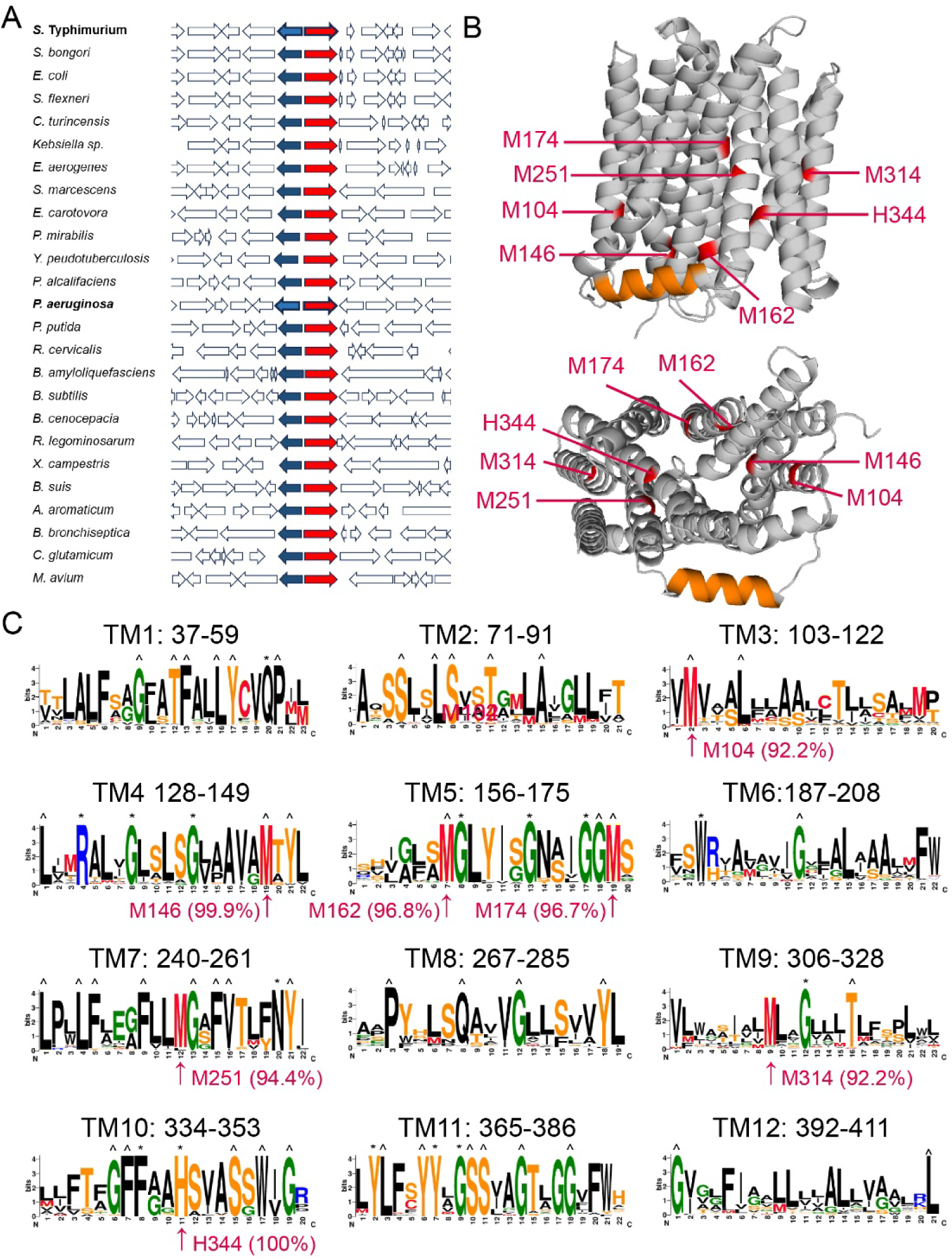
Evolutionary Conservation and Structural Features of the Putative Bacterial Cu-Transport Protein CuiT. **(A)** Conservation of *cuiT* (red) and its putative regulatory gene (blue) across phylogenetically diverse bacterial species. Shown are representative chromosomal regions from bacteria harboring *cuiT* homologs, illustrating conserved gene synteny. Analysis was performed using the SEED Viewer platform. A single representative strain of *S. enterica* or *Salmonella bongori* is shown; all sequenced *Salmonella* genomes exhibit an identical chromosomal arrangement at this locus. **(B)** AlphaFold-predicted three-dimensional model of CuiT. Membrane and cytoplasmic side views. Conserved residues possibly participating of binding/transport are highlighted in red. The orange helix is a hypothetical chaperone docking site. **(C)** Predicted transmembrane segments of *Salmonella* CuiT. Sequences logos represent the conservation of individual residues in the 792 sequences analyzed. Annotations represent: ^ 95-99.9% and * 100% conservation and ↑ points to conserved residues likely facing the cation transport path or Cu-binding sites.

The AlphaFold 3-predicted three-dimensional structure reveals that CuiT contains 12 transmembrane α**-**helices of archetypical MFS transporter (**Figure 1B**) (42). The hypothetical structure also shows a clear division between the outward-facing (extracellular/periplasmic) and inward-facing (cytoplasmic) regions, suggesting an alternating access mechanism where conformational changes allow Cu uptake and release into the cytoplasm. The structure contains a singular cytoplasmic transversal helix (orange in **Figure 1B**) that resembles a similar structure of P-type Cu-ATPases that participates in Cu-chaperone docking (9, 43). This docking is key for the Cu transfer among transporters and chaperones (9). Sequence conservation can be applied to identify residues that are crucial for protein functions such as catalytic activity, ligand binding, or maintaining the stability of the folded structure (44). Analysis of the sequences of putative transmembrane segments shows highly conserved His, Met, and potentially Cys residues (**Figure 1C**). These amino acids are known to participate in Cu binding and coordination in other transporters, facilitating the selective movement of the ion across the membrane (1, 4, 10, 45–49). In particular, His and Met residues are enriched in TM1, TM4, TM6, TM9, TM10, and TM11 (**Figure 1C**). Notably, Met residues (MET104, MET146, MET162, MET174, MET251, MET314) and a His residue (HIS344) are positioned along a putative transport path, reinforcing the idea of their potential role in Cu /Cu^2+^ coordination and translocation (1, 4, 48) (**Figure 1B**).

### CuiT enhances Cu tolerance in *Salmonella*

Bacterial Cu tolerance can be used as an early simple approach to explore the participation of a gene in Cu homeostasis. To this end, the growth rate of wild-type (WT), Δ*cuiT* mutant and complemented strains, and cells overexpressing *cuiT* under control of a P*lac* promoter were analyzed under increasing CuSO_4_ concentrations. Growth curves in LB broth show that in the absence of CuSO_4_ (0 mM), all strains exhibited comparable growth rates, indicating that the deletion of CuiT does not affect basal growth (**Figure 2A**). However, complementation of Δ*cuiT* (Δ*cuiT*/pCuiT) and overexpression of CuiT (WT/pCuiT) resulted in slightly impaired growth compared to WT at 4 mM CuSO_4_, suggesting that the transporter increases the entrance of Cu and therefore these strains showed a slightly lower metal tolerance (**Figure 2A**). At 6 mM CuSO_4_, all strains exhibited growth inhibition.

**Figure 2.**
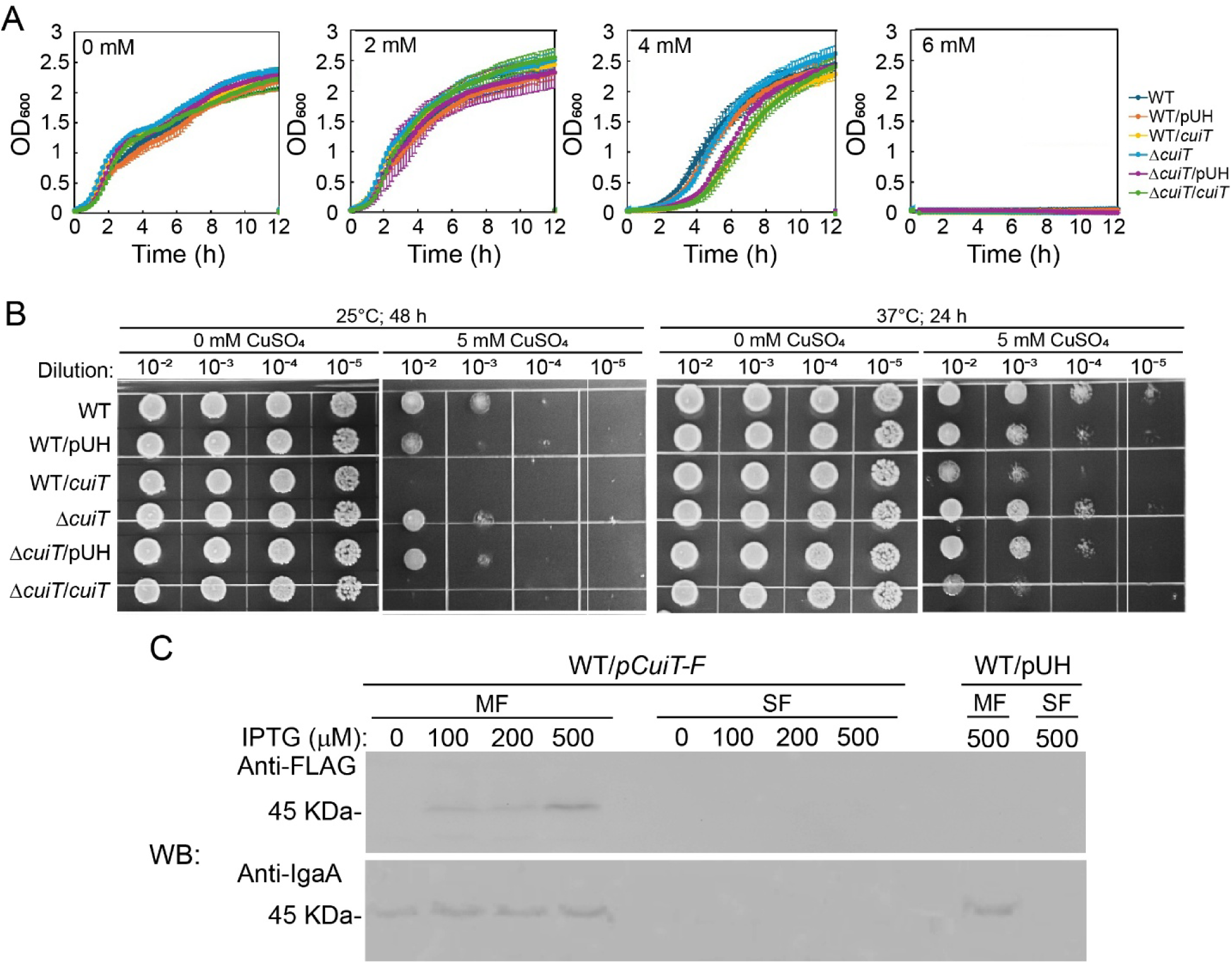
CuiT confers Cu sensitivity in *Salmonella enterica* serovar Typhimurium. **(A)** Growth analysis of wild-type *S. enterica* WT), *cuiT* overexpressing WT (WT/pCuiT), *cuiT* deletion mutant (Δ*cuiT*), complemented (Δ*cuiT*/pCuiT) and empty vector controls (WT/pUH and Δ*cuiT*/pUH) strains, cultured in LB medium supplemented with increasing concentrations of CuSO_4_ (0-6 mM). **(B)** Cu sensitivity of spot-plated serial dilutions of strains shown in (A) onto LB agar with or without 4 mM CuSO_4_. **(C)** Recombinant CuiT protein localizes to the membrane fraction. WT cells harboring pCuiT-3×FLAG (pCuiT-F) or the empty vector (pUH) were grown, subjected to subcellular fractionation and analyzed by SDS-PAGE and immunoblotting using anti-FLAG (α-FLAG) or anti-IgaA (α-IgaA) antibodies, as indicated. The inner membrane protein IgaA, was included as a loading control and membrane marker.

Tolerance in solid media was performed to further validate these observations (**Figure 2B**). Interestingly, the overexpressing strain WT/pCuiT and the complemented strain Δ*cuiT*/pCuiT exhibited an even clearer phenotype with a markedly reduced viability on LB agar plates supplemented with 5 mM CuSO_4_, whereas WT, Δ*cuiT* and their empty vector control (WT/pUH, Δ*cuiT*/pUH) showed similar growth even when exposed to high CuSO_4_. These Cu sensitivity phenotypes were consistent at both 25°C and 37°C. Western blot analyses of membrane and soluble cellular fractions confirm that IPTG induced CuiT expression resulted in the membrane-localized transporter, in a similar manner that a known membrane protein, the intracellular growth attenuator A (IgaA) **Figure 2C**). In summary, the data is consistent with CuiT enabling the entrance of Cu with a resulting loss of tolerance to the metal.

### Expression of *cuiT* gene upon Cu stress

We reported that *cuiT* expression is repressed in *Pseudomonas* upon Cu stress (31). A similar phenomenon was observed in *Salmonella* when the relative *cuiT* expression was assessed in cells exposed to increasing concentrations of CuSO_4_ for 5 min (**Figure 3A**; black bars) and 60 min (**Figure 3A**; gray bars). This is, *cuiT* expression was repressed by increased intracellular Cu. Testing the hypothesis that deletion of *cuiT* would reduce intracellular Cu, the expression of key Cu homeostasis genes (*copA, cueP, cueO, golB,* and *scsC*) in WT and Δ*cuiT* mutant strains was analyzed after 5 min of exposure to 4 mM CuSO_4_ (**Figure 3B**). As expected, the Δ*cuiT* mutant exhibited a marked decrease in expression across all analyzed genes, with the most significant reduction observed for Cu chaperones *golB* and *cueP*. These findings are congruent with CuiT mediating cellular Cu influx.

**Figure 3.**
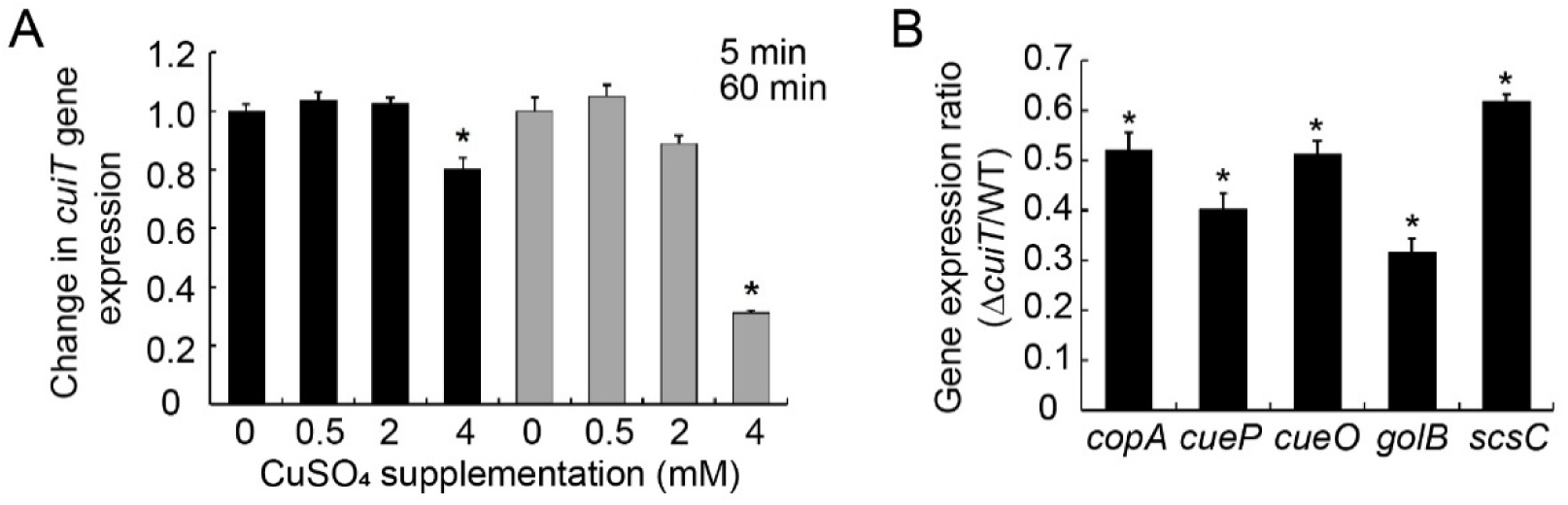
CuiT expression is regulated by Cu and affects the expression of Cu-responsive genes in *Salmonella.* **(A)** Cu-responsive *cuiT* gene expression in wild-type (WT) cells following exposure to increasing concentrations of CuSO_4_ for 5 min (black bars) or 60 min (gray bars). Data were normalized to the housekeeping gene *rnpB* and expressed relative to time 0. Data are the mean ± S.E. of three independent experiments (n=3). Statistical significance was determined using an unpaired Student’s t test (*p<0.05). **(B)** Cu-responsive expression of Cu homeostasis genes in Δ*cuiT* strains relative to that in WT, following treatment with 1 mM CuSO_4_ for 5 min. Data were normalized to the housekeeping gene *rnpB* and expressed relative to time 0. Data are the mean ± S.E. of three independent experiments (n=3). Statistical significance was determined using an unpaired Student’s t test (p<0.05).

### CuiT modulates intracellular Cu and its uptake kinetics in *Salmonella*

Seeking a direct demonstration of the role of CuiT as a Cu importer, the Cu contents in WT and Δ*cuiT* mutant strains were measured. Both *WT* and Δ*cuiT* strains accumulated Cu in a dose-dependent manner under increasing CuSO concentrations (0-4 mM). However, at 4 mM CuSO, the Δ*cuiT* mutant exhibited significantly lower intracellular Cu levels compared to *WT* (**Figure 4A**). Toward further confirming the role of CuiT, the Cu levels in complemented and CuiT-overexpressing strains were measured (**Figure 4B**). As shown, Δ*cuiT* and Δ*cuiT*/pUH strains showed significantly lower Cu content. Interestingly, the complementation of Δ*cuiT* with a plasmid expressing CuiT (Δ*cuiT*/pCuiT) restored Cu accumulation to levels comparable to the WT, while CuiT overexpression (WT/pCuiT) resulted in a marked increase in intracellular Cu. These results indicate that CuiT plays a critical role in Cu import.

**Figure 4.**
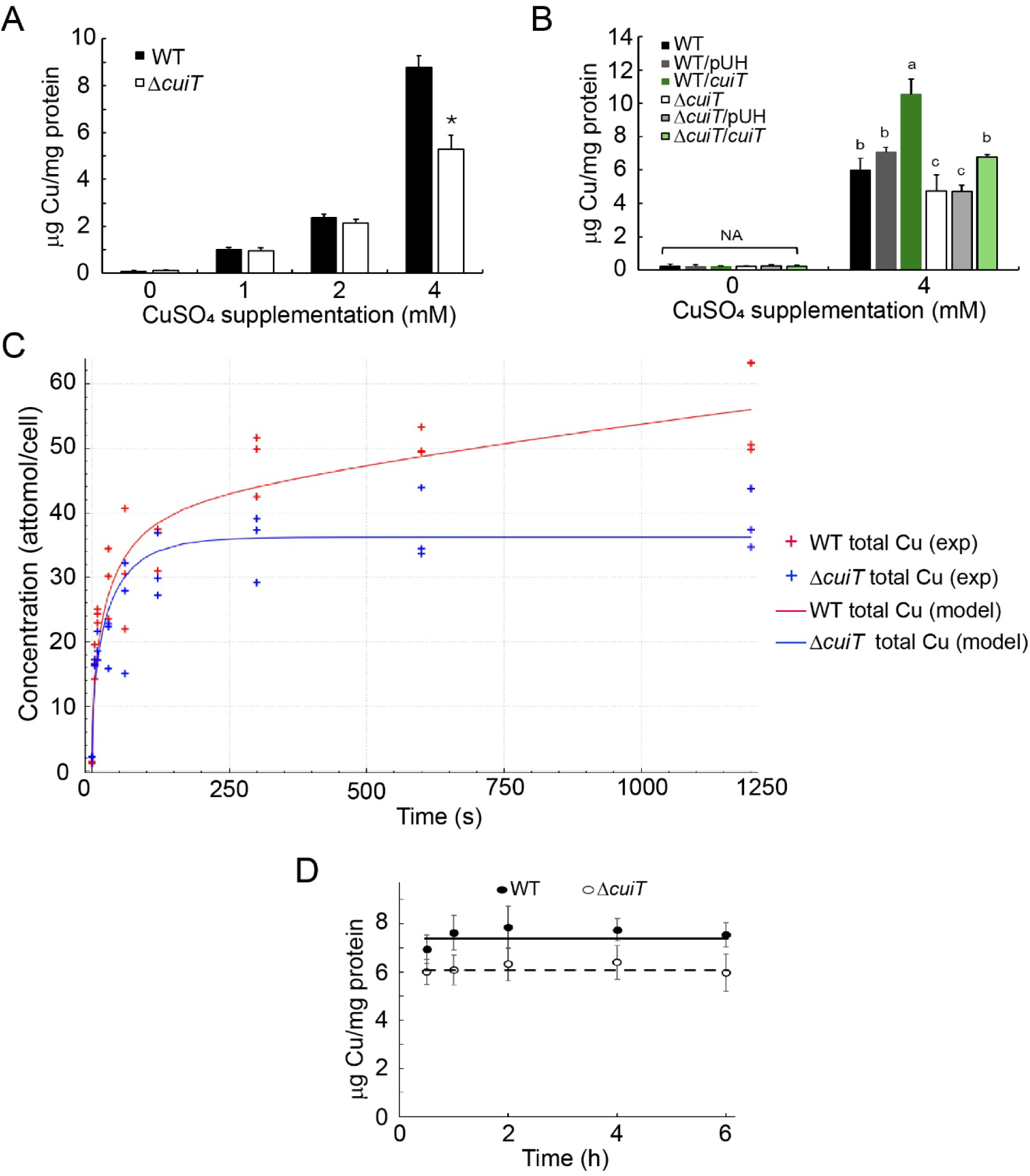
CuiT regulates Cu uptake kinetics and intracellular Cu accumulation in *Salmonella.* **(A)** Intracellular Cu accumulation in WT and Δ*cuiT* mutant strains following exposure to increasing concentrations of CuSO_4_ for 1 h. **(B)** Intracellular Cu levels in WT, Δ*cuiT*, complemented (Δ*cuiT*/pCuiT), and *cuiT*-overexpressing strains grown in the absence or presence of 4 mM CuSO_4_ for 1 h. Data are mean ± S.E. of three independent experiments. Statistical significance was determined using an unpaired two-tailed Student’s t test (p < 0.05). **(C)** Cu uptake kinetics in WT and Δ*cuiT* mutant strains following exposure to 4 mM CuSO_4_. Red and blue lines result of curve fitting using the computational compartmental distribution model described in Figure 5. Fitting parameters are listed in **Table 1**. Cu-uptake into the Δ*cuiT* strain was modeled making the uptake through CuiT null (*v*CuiT=0). Data are the mean ± S.E. of three independent experiments (n=3). **(D)** The intracellular Cu levels of WT and Δ*cuiT* mutant strain measured in an extended timeframe in the presence of 4 mM CuSO_4_. Data are the mean ± S.E. of two independent experiments.

**Table 1.** The fit parameter values for Cu uptake kinetics. Comparison between *Pseudomonas* and *Salmonella* models.

| Parameter | *Pseudomonas* | Parameter | *Salmonella* | Unit |
| --- | --- | --- | --- | --- |
| $v_{\text{CcoA}}$ | 4.43 | $v_{\text{CuiT}}$ | 380 | mM/min |
| $K_m_{\text{CcoA}}$ | 0.00950 | $K_m_{\text{CuiT}}$ | 0.0149 | mM |
| $v_{\text{OprC}}$ | 6.84 | $v_{\text{I-Porin}}$ | 210 | mM/min |
| $K_m_{\text{OprC}}$ | 1.66 | $K_m_{\text{I-Porin}}$ | 0.252 | mM |
| $v_{\text{CopA1}}$ | 0.713 | $v_{\text{CopA/GolT}}$ | 100 | mM/min |
| $K_m_{\text{CopA1}}$ | 0.000476 | $K_m_{\text{CopA/GolT}}$ | 0.000354 | mM |
| $v_{\text{PcoB}}$ | 3.68 | $v_{\text{E-Porin}}$ | 200 | mM |
| $K_m_{\text{PcoB}}$ | 0.390 | $K_m_{\text{E-Porin}}$ | 0.605 | mM |
| $k_{\text{ds}}\text{CuCytCP}$ | 98.6 | $k_{\text{ds}}\text{CuCytCP}$ | 187 | 1/min |
| $k_{\text{ds}}\text{CuPPCP}$ | 0.00126 | $k_{\text{ds}}\text{CuPPCP}$ | 0.264 | 1/min |
| $K_d\text{CytCP}$ | 0.000218 | $K_d\text{CytCP}$ | 0.00220 | mM |
| $K_d\text{PPCP}$ | $2.13 \times 10^{-7}$ | $K_d\text{PPCP}$ | 0.00525 | mM |
| $[\text{CuCytCP}]$ | 0.00892 | $[\text{CuCytCP}]$ | 0.0178 | mM |
| $[\text{CuPPCP}]$ | 0.419 | $[\text{CuPPCP}]$ | 0.837 | mM |

\* Previously reported values obtained for *Pseudomonas aeruginosa* (30).
$v$ are $V_{\text{max}}$ and $K_m$ the Michaelis constant for the corresponding transporter.
$k_{\text{ds}}$ are dissociation constants for Cu binding to the corresponding chaperone
$K_d$ are the equilibrium constants for the metal transfer reaction between the chaperone and the corresponding cuproprotein.
$[\text{CuCytPP}]$ and $[\text{CuPPCP}]$ are the size of the corresponding cuproprotein pools.

To further examine the role of CuiT in Cu transport, we monitored the kinetic of Cu uptake in the WT and the Δ*cuiT* mutant strain exposed to 4 mM CuSO_4_. The WT cells rapidly accumulated Cu, reaching steady-state level within 15-20 min (**Figure 4C**). In contrast, the Δ*cuiT* strain exhibited a slower Cu uptake rate with reduced overall accumulation once steady state is reached (**Figure 4C**). Interestingly, the distinct steady state Cu levels can be maintained (**Figure 4D**) and, as shown, it does not visibly affect the cell viability (**Figure 2A**). We have previously demonstrated that the whole cell Cu uptake kinetics and the steady state metals levels can be modeled using compartmental analysis (**Figure 5**) (30). In this approach, the transmembrane transporters are defined by Michaelis-Menten kinetics parameters (V_max_ and K_m_), while Cu transfer among chaperones and cuproproteins are described by affinity (k_ds_)and equilibrium (K_d_) constant (**Table 1**). These parameters are integrated in a full equation that defines Cu content versus time. We initiate the fitting of our data using the model designed for *Pseudomonas aeruginosa.* However, this model performed poorly, *i.e*., the fitting curve did not match the experimental data. We hypothesize that the issue could be due to the inclusion of a direct transfer of Cu from the cytoplasm to the extracellular media via a CusABCF type system, since this transporter is absent in *Salmonella.* Removal of CusABCF as shown in **Figure 5** allowed fitting the curves, as well as providing support to the model strength. Further confirming the validity of the approach and the role of CuiT as the plasma membrane Cu influx transporter, the Δ*cuiT* strain data could be fitted by simply making the plasma membrane influx null (*v*CuiT = 0).

**Figure 5.**
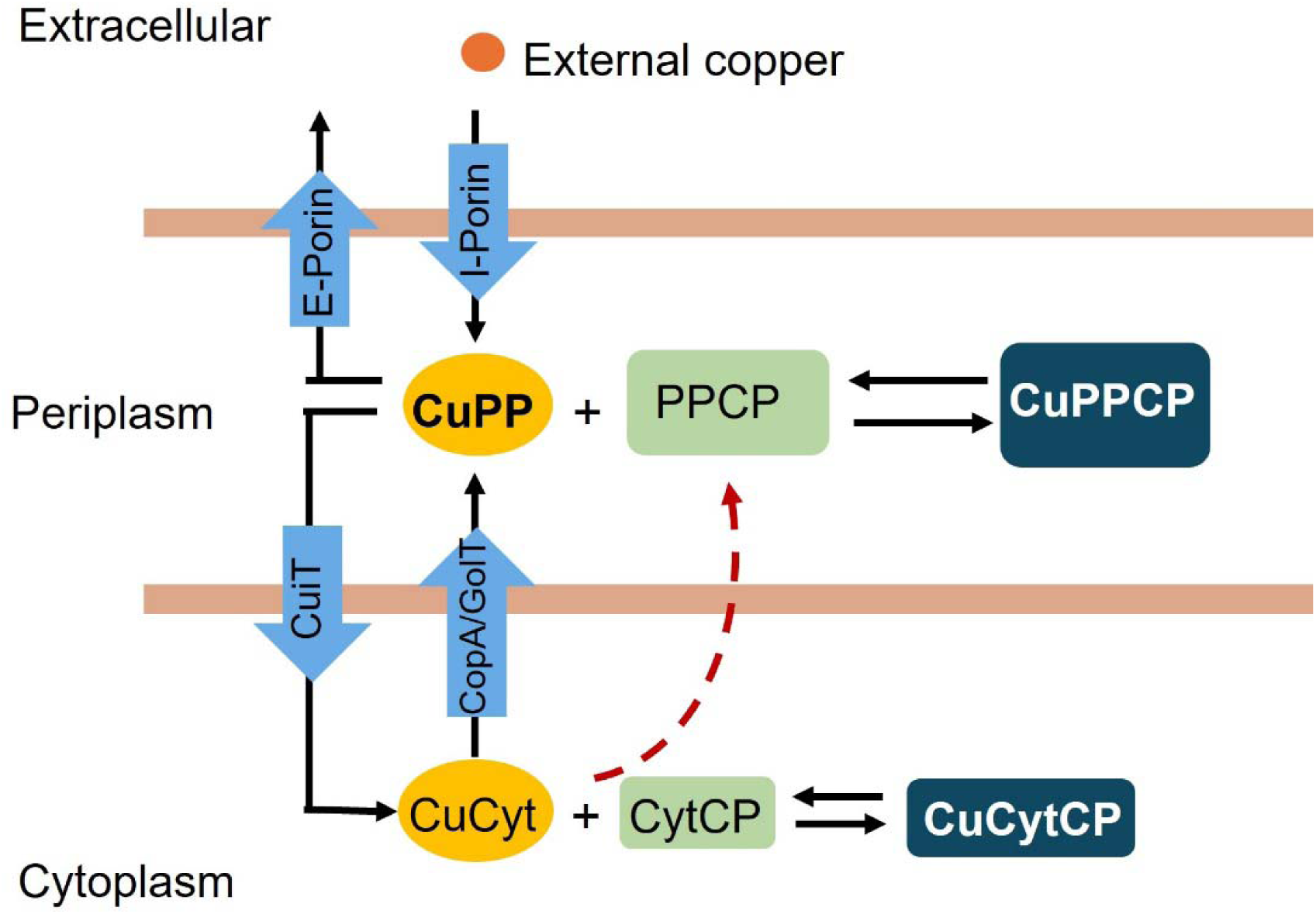
Model of the Cu homeostasis network in *Salmonella*. Data of Cu uptake kinetics was analyzed with a computational compartmental distribution model (5, 30). Our previously reported model describing the Cu uptake kinetics into *Pseudomo*nas aeruginosa was modified by removing equations describing an ABCF transporter absent in Salmonella (50). The model assumes the presence of outer membrane porins for influx and efflux. The plasma membrane contains influx (CuiT) and efflux (Cu-ATPases) systems. Various soluble Cu chaperones (CuPP and CuCyt) that carry Cu to target cuproproteins and membrane transporters. The model requires the upregulation (dotted line) of the periplasmic Cu pool. As a consequence of the predicted high affinity binding to all systems the model implies the absence of free Cu.

**Table 1** lists the fitting parameters for *Salmonella* and includes for reference the previously reported equivalent values observed in *Pseudomonas*. *Salmonella* showed a faster Cu uptake rate (*v*CuiT) albeit with lower affinity compared to *Pseudomonas*. Additionally, *Salmonella* exhibited higher parameter values for Porin-mediated uptake. This was logically parallelled by an increase in the efflux rates by P-type ATPases and E-Porins. This means that the *Salmonella* Cu pool has a higher turnover rate. Also in agreement are the relatively higher Cu pool in *Salmonella* with and observed slightly higher K_eq_ for the metal transfer. In spite of these kinetics differences, the Cu chaperone affinities (k_ds_) are similar in both systems, *Salmonella* and *Pseudomonas,* indicating that they operate in the absence of free/hydrated metal as expected.

## DISCUSSION

### CuiT-dependent Cu stress phenotypes reveal a critical role in Cu acquisition

Cu homeostasis requires a precise balance between acquisition and detoxification, as both Cu deficiency and excess are detrimental to bacterial physiology (1, 2, 5, 7). In this study, we identify CuiT as a key determinant of Cu stress phenotypes in *S. enterica*. Unlike canonical Cu efflux systems, whose deletion leads to dramatic sensitivity to the ion and growth defects (1, 4, 5, 12), loss of CuiT did not impair basal growth or gross Cu tolerance under moderate Cu stress. Instead, CuiT overexpression or complementation under Cu stress conditions led to heightened Cu sensitivity, as evidenced by impaired growth and reduced viability in spot assays. These phenotypes are consistent with CuiT functioning primarily in Cu acquisition rather than export. The observed sensitivity associated with CuiT overexpression likely reflects excessive intracellular Cu accumulation beyond the buffering capacity of the cellular Cu-handling network. This interpretation is supported by the elevated intracellular Cu levels detected in the CuiT-overexpressing strain compared to WT cells. Such Cu overload can exacerbate oxidative damage, thiol oxidation, and metal mismetallation, processes well known to underlie Cu toxicity (5, 7, 8). Thus, while CuiT-mediated Cu import is beneficial under Cu-limiting or physiological conditions, its dysregulation becomes detrimental under high Cu stress, emphasizing the necessity for tight control of Cu influx pathways.

Notably, the Cu stress phenotypes associated with CuiT differ from those reported for *Pseudomonas*, where deletion of the CuiT homolog conferred enhanced Cu tolerance in solid media (31). These differences likely reflect species-specific adaptations in Cu handling networks, particularly given that *Salmonella* lacks the CusCFBA efflux system and relies heavily on CueP and other periplasmic mechanisms to buffer Cu stress (39, 50). Deletion of *cuiT* resulted in a broad attenuation of Cu-responsive gene expression, including *copA, cueP, cueO, golB,* and *scsC*, indicating that this transporter is required for full activation of the Cu homeostasis network under Cu stress. The most pronounced reductions were observed for the Cu chaperones *golB* and *cueP*, suggesting that impaired Cu import in the Δ*cuiT* mutant limits intracellular Cu availability necessary to activate CueR- and GolS-dependent transcriptional responses. These findings support a model in which CuiT-mediated Cu uptake functions upstream of Cu-sensing regulators, ensuring sufficient intracellular Cu to trigger protective and adaptive gene expression programs while coordinating metal export, chaperoning, and redox stress resistance pathways (5, 37, 51). Thus, CuiT emerges as a contributor to Cu stress phenotypes in *Salmonella*, functioning at the intersection of Cu acquisition and toxicity.

### Integration of CuiT into the *Salmonella* Cu homeostasis network

Multiple lines of evidence from this study establish CuiT as a central component of the *Salmonella* Cu homeostasis network. Comparative genomic analyses revealed that CuiT is highly conserved across *Enterobacteriaceae* and present in diverse bacterial taxa, often within conserved genomic neighborhoods (41). AlphaFold 3 structural modeling predicts that CuiT presents a 12-transmembrane-helix topology, characteristic of major facilitator superfamily transporters, forming a central channel compatible with an alternating-access transport mechanism (42). CuiT proposed topology closely resembles the established Cu importers CcoA in *R. capsulatus* and YcnJ in *B. subtilis* (15, 16, 52). Consistently, conserved His, Met, and Cys residues are enriched within key transmembrane segments and are strategically positioned along the predicted transport pathway, foreseen from established principles of Cu coordination and translocation (45–48). The conservation of both sequence and structural features across bacterial lineages indicates that CuiT represents an evolutionarily maintained solution for Cu acquisition, reinforcing its functional importance in bacterial Cu homeostasis. CuiT evolutionary conservation suggests a broadly preserved role in bacterial Cu physiology, analogous to other characterized Cu importers such as CcoA and YcnJ (15–17).

Functionally, deletion of CuiT resulted in significantly reduced intracellular Cu accumulation under Cu stress, while complementation restored Cu levels to those observed in WT cells. Moreover, *cuiT* overexpression led to Cu accumulation far exceeding control levels, directly implicating the protein in Cu import or retention. Kinetic analyses further demonstrated that CuiT is required for rapid Cu uptake during the early phases of Cu exposure, with the Δ*cuiT* mutant displaying delayed and diminished Cu accumulation. These findings indicate that CuiT contributes to both the rate and magnitude of intracellular Cu acquisition. Strikingly, kinetic modeling revealed that CuiT-mediated Cu uptake in *Salmonella* occurs at a substantially higher rate than CcoA-mediated uptake in *P. aeruginosa*, accompanied by a large intracellular Cu pool. This enhanced uptake capacity may reflect the adaptation capacity of *Salmonella* to host-associated environments, where Cu is deployed as an antimicrobial weapon during the oxidative burst (6, 31, 50). This divergence likely reflects adaptation to host-associated environments, where *Salmonella* encounters fluctuating and often elevated Cu concentrations as part of the innate immune response (6). In contrast, Cu importers in environmental bacteria such as *R. capsulatus* and *B. subtilis* may be optimized for basal Cu acquisition to support cuproenzyme assembly rather than rapid Cu influx under stress. Thus, CuiT appears to represent an evolutionarily conserved yet functionally specialized Cu importer, tuned to meet the distinct Cu demands imposed by pathogenic lifestyles.

Efficient Cu acquisition may be essential not only for cuproenzyme metalation but also for sustaining Cu-dependent defense systems, such as SodCI and SodCII, whose activity is supported by CueP-mediated Cu delivery (37, 51). Consistent with this model, loss of CuiT resulted in reduced expression of multiple Cu-responsive genes, including *copA, cueP, cueO, golB*, and *scsC*. This coordinated downregulation suggests that CuiT-dependent Cu import is necessary to activate Cu-sensing regulatory circuits, likely through CueR- and GolS-mediated transcriptional responses (37). Thus, CuiT acts upstream of transcriptional and enzymatic components of Cu homeostasis, positioning it as a key gatekeeper of intracellular Cu availability in *Salmonella*.

### Cu-dependent control of CuiT expression and transport activity

The expression and activity of Cu import systems must be tightly regulated to prevent Cu toxicity, and our data indicate that CuiT is subject to Cu-responsive regulation. Exposure to high Cu concentrations led to a rapid and sustained decrease in *cuiT* transcript levels, suggesting that CuiT expression is actively repressed under Cu stress. This regulatory behavior mirrors that of other Cu importers, such as *oprC* in *Pseudomonas*, whose expression is downregulated in response to elevated Cu levels (26, 27, 31). Such repression likely serves as a protective mechanism to limit further Cu influx when intracellular Cu is already abundant. The repression of Cu-responsive genes in the Δ*cuiT* mutant suggests that CuiT-mediated Cu import is required to reach intracellular Cu thresholds necessary for activation of Cu-sensing regulators such as CueR and GolS (37). In this context, CuiT does not merely function as a passive transporter but plays an active role in shaping Cu-dependent regulatory responses. Downstream, CueP and the ScsABCD system may buffer and redistribute imported Cu, linking CuiT activity to periplasmic Cu handling, redox balance, and stress resistance in the absence of the CusCFBA efflux system (6, 37, 51). Thus, the integration of CuiT-mediated import with transcriptional control ensures that Cu uptake is dynamically tuned to environmental availability and cellular demand.

Collectively, our findings identify CuiT as an evolutionarily conserved Cu importer that plays a critical role in *Salmonella* Cu homeostasis. CuiT facilitates rapid Cu uptake, supports activation of Cu-responsive gene expression, and must be tightly regulated to avoid Cu toxicity. Given the central role of Cu in host-pathogen interactions and innate immune defense (5, 53), CuiT likely contributes to *Salmonella* fitness during infection. These results expand the current framework of bacterial Cu homeostasis by highlighting Cu import, rather than efflux alone, as a crucial determinant of Cu stress adaptation in pathogenic bacteria. CuiT and its associated regulatory networks represent promising antimicrobial targets that exploit vulnerabilities in bacterial Cu homeostasis while minimizing effects on host metal metabolism.

## MATERIALS AND METHODS

### Bacterial strains and culture conditions

Bacterial strains and plasmids applied in this study are listed in **Supplemental Table 1**. Bacterial cells were routinely cultured at 37°C in Luria-Bertani (LB) broth or on LB-agar plates, except when indicated. When necessary, the culture medium was supplemented with chloramphenicol at 34 µg/ml and/or ampicillin at 100 µg/ml. When indicated 0.5 µM isopropyl β-D-1-thiogalactopyranoside (IPTG) was added to express CuiT from a plasmid. Cu sulfate salt used were of ACS analytical grade ≥98.0% purity.

### Mutant strain construct

The ATCC 14028s Δ*cuiT* derivative was generated by Lambda Red-mediated recombination using the primers listed in **Supplemental Table 2** and previously described protocols (40). Then, the *cuiT*::*Cm^R^* modification was moved to non-modified 14028s strain by P22-mediated transduction. DNA fragments and plasmids were introduced into bacterial cells by electroporation. All constructs were verified by DNA sequencing.

For generation of pCuiT and pCuiT-F plasmids, the *cuiT* (STM1486) open reading frame was amplified by PCR from the chromosome of the WT 14028s strain using the oligonucleotides listed in Supplemental Table S2. The product containing *cuiT* was digested with *BamH*I/*Hind*III and cloned into the pUH21-2lacI*^q^* plasmid digested with these enzymes. The plasmids were initially introduced into competent *E. coli* Top10 cells. Positive clones were verified by colony PCR and DNA sequencing.

### Western blot analysis

Western blot analysis of 3xFLAG-tagged CuiT was done as described previously (5). Briefly, 14028s cells carrying either the empty vector or the pCuiT-3xFlag plasmid were grown overnight at 37°C in LB broth supplemented with 100 mg/ml ampicillin. When the cultures reached OD_600_ ∼0.5, 0.2 mM IPTG was added and cell incubated overnight. Cells were harvested by centrifugation at 6000 rpm for 10 min, washed and resuspended in 1/10 of the original volume of 50 mM Tris-HCl pH 8, 5mM EDTA, 10% Glycerol solution supplemented with 1 mM phenylmethylsulfonyl fluoride (PMSF) and 50 mM β-mercaptoethanol. Cell suspensions were sonicated on ice centrifuged at 6000 rpm for 10 min at 4°C to eliminate non-disrupted bacteria. The supernatant was subjected to ultracentrifugation for 60 min at 45,000 rpm at 4°C to obtain soluble and insoluble (membrane) fractions. The pellets were solubilized in buffer containing 50 mM Tris-HCl pH 8, 5mM EDTA, 10% Glycerol and 2% SDS. Pierce BCA Protein Assay Kit (Thermo Scientific) was used to determine protein concentration in total, soluble or insoluble extracts following the manufacturer’s protocols. Aliquots containing 20 µg of protein were analyzed in 15% (w/v) SDS polyacrylamide gels. Two gels processed in parallel were transferred to nitrocellulose membranes (Amersham Protran Premium 0.45 µm NC, Cytiva) for either CuiT-3xFLAG or IgaA immunodetection, using mouse anti-FLAG monoclonal (ab125243, Abcam) or rabbit anti-IgaA (ab97216, Abcam) polyclonal antibodies, respectively. Blotted membranes were incubated with the corresponding secondary antibodies conjugated with horseradish peroxidase (HRP; AP106P and 12-349, Sigma). Immunoreactive bands were revealed using the Pierce™ ECL Western Blotting Substrate” (Thermo Fisher Scientific) and registered in a ChemiDoc™ XRS Imaging System (Bio-Rad).

### Cu sensitivity assay

Bacterial cultures at the mid-log phase (OD_600_=0.5-0.6) were diluted in LB medium to achieve an initial OD_600_=0.05. Following dilution, the cultures were supplemented with varying concentrations of CuSO_4_, ranging from 0 to 6 mM. Cell growth in 0.2 ml liquid medium was tracked over 16 h at 37°C with continuous agitation using an Epoch 2 microplate spectrophotometer (BioTek) (4). OD_600_ readings were taken regularly to evaluate bacterial growth dynamics. Cu tolerance in solid media was assessed using LB-agar plates supplemented with increasing concentrations of CuSO_4_. Mid-log phase bacterial cultures (OD_600_ = 0.5-0.6) were diluted to an OD_600_ = 0.1 and further serially diluted down to an OD_600_ of 10^-6^ in LB. Subsequently, 10 µl of each dilution was spot plated onto the LB-agar plates. The plates were incubated at either 25°C or 37°C to allow colony growth (54).

### Gene expression analysis

*Salmonella* WT and Δ*cuiT* strains in mid-log phase were grown during 5 or 60 min at 37°C in LB medium supplemented with the indicated CuSO_4_ with 0, 0.5, 2 and 4 mM of CuSO_4_ at 37°C. Aliquots of 0.5 mL were stabilized with RNA protect Bacteria Reagent (Qiagen), RNA was isolated with RNeasy Mini Kit (Qiagen), treated with DNase (Qiagen). RNA (500 ng) was used for cDNA synthesis using the iScript™ gDNA Clear cDNA Synthesis Kit (Bio-Rad). Quantitative PCR (qPCR) were carried out using 25 ng of cDNA in a LightCycler® 96 System (Roche) with AzuraView™ GreenFast qPCR Blue Mix LR (Azura Genomics) in a 20 μl final volume, using 0.8 μM of each primer. *rnpB* was used as the reference gene. Gene-specific primers are listed in **Supplementary Table 2** (54, 55).

### Whole-cell Cu uptake assays

Cu levels were determined by atomic absorption spectroscopy (AAS), as described (31). Briefly, bacterial cells in mid-log phase were incubated in LB medium supplemented with varying concentrations of CuSO_4_ during the times indicated in the figures. Subsequently, a fraction of the cell suspension (100 µl) was reserved for protein quantification using the Bradford method (56), while the remaining portion (1 ml) was passed through LabExact Mixed Cellulose Ester (MCE) membrane filters (0.45 µm pore size, 25 mm diameter, Global Industrial, Model #: WBB2684613) under vacuum conditions. The filters were subsequently washed twice with 150 mM NaCl (30 ml) under vacuum. Following filtration and washing, the filters were air-dried and transferred into cap crewed Micrewtubes (Simport #T339). The cells with the filters were mineralized using fuming HNO_3_ (trace metal grade; 400 µl) at 80°C for 60 min. After cooling, the mineralized samples were treated with 10 M H_2_O_2_ (200 µl) and ddH_2_O (400 µl) for an additional 60 min at room temperature. Resulting samples were analyzed by AAS.

### Kinetic analysis

Kinetic analysis and parameter fitting were carried out with the software COPASI (v. 4.45; (57)), using a kinetic model based on that previously described for *Pseudomonas* (30) as described in the legend of Figure 5.

### Bioinformatics analysis

NCBI Reference Sequence WP_000091796.1 was used to Blast the proteins with NCBI RefSeq Select proteins (**Supplemental Table S3**). The resulting 1000 sequences with higher similarity were screened removing redundant and incomplete sequences. The selected 792 sequences were aligned in MuscleWS. The alignment was visualized in Jalview Version 2.11.3.2 (58) and the sequence logos were generated in WebLogo (59, 60). The structure of the protein was predicted in AlphaFold 3 (42). The structure was visualized and modeled in ChimeraX-1.7.1 (61–63).

## Supporting information

Zhao Supplemental table 3

## ACKNOWLEDGEMENTS

We thank Julián Mendoza, who conducted some preliminary experiments with the *cuiT* mutant. This work was supported by the National Institutes of Health (R01A1150784 to J. M. A.). T.P.-B. was supported by Wesleyan University institutional funds. P.M. was supported by the National Institutes of Health (R24GM137787). A.A.E.M. is a CONICET fellow. S.K.C and F.C.S. are career investigators of CONICET. F.C.S. is also a Career Investigator of Consejo de Investigaciones de la Universidad Nacional de Rosario.

## AUTHOR CONTRIBUTIONS

Zhenzhen Zhao: Data acquisition, curation, analysis, validation and visualization; manuscript-first draft, review and editing.

Karla F. Díaz Rodríguez: Data acquisition, curation, analysis, validation and visualization; manuscript-review and editing.

Andrea A. E. Méndez: Preparation of mutant strains, curation, analysis, and validation; manuscript-review and editing.

Lisandro M. Somers: Preparation of mutant strains curation, analysis, and validation; manuscript-review and editing.

Pedro Mendes: Computational analysis of Cu uptake data, validation and visualization; manuscript-review and editing.

Fernando Soncini: genomic analysis, investigation, and visualization; Manuscript-review and editing, supervision.

Susana K. Checa: Manuscript-review and editing, supervision.

Teresita Padilla-Benavides: Writing first draft, review and editing; funding acquisition.

José M. Argüello: Experimental design, methodology, data curation and analysis, manuscript first draft, review and editing, funding acquisition, supervision.

---

**Supplemental Table S1.** Bacterial strains and plasmids used in this study.

| Strain | Relevant genotype or properties | Refs. or source |
| --- | --- | --- |
| 14028s | Wild type | ATCC®-14028™ |
| $\Delta cuiT$ | Mutation in CuiT (Cm<sup>R</sup>) | This work |
| $\Delta cuiT$/pCuiT | cuiT complemented strain, $\Delta cuiT$ transformed with pUH::CuiT (Cm<sup>R</sup>+Amp<sup>R</sup>) | This work |
| WT/CuiT | cuiT overexpress strain, Wild type transformed with pUH::CuiT (Amp<sup>R</sup>) | This work |
| WT/pUH | Wild type transformed with empty vector pUH21.2-*lacI*<sup>q</sup> vector (Amp<sup>R</sup>) | This work |
| $\Delta cuiT$/pUH | $\Delta cuiT$ transformed with empty vector pUH21.2-*lacI*<sup>q</sup> vector (Cm<sup>R</sup>+Amp<sup>R</sup>) | This work |
| Plasmid | Relevant genotype or properties | Refs. or source |
| pKD46 | oriR<sub>pSC101</sub>ts P<sub>araB</sub> *exo-bet-gam* Amp<sup>R</sup> | (64) |
| pKD3 | oriR6K FRT-Cm<sup>R</sup>-FRT Amp<sup>R</sup> | (64) |
| pUH21–2 *lacI*<sup>q</sup> | reppMB1 Ap<sup>R</sup> *lacI*<sup>q</sup> | (65) |
| pPB1496 (pCuiT) | pUH21–2 *lacI*<sup>q</sup> derived plasmid carrying *cuiT* | This work |
| pPB1498 (pCuiT-F) | pUH21–2 *lacI*<sup>q</sup> derived plasmid carrying *cuiT*-3xFLAG | This work |

**Supplemental Table S2.**
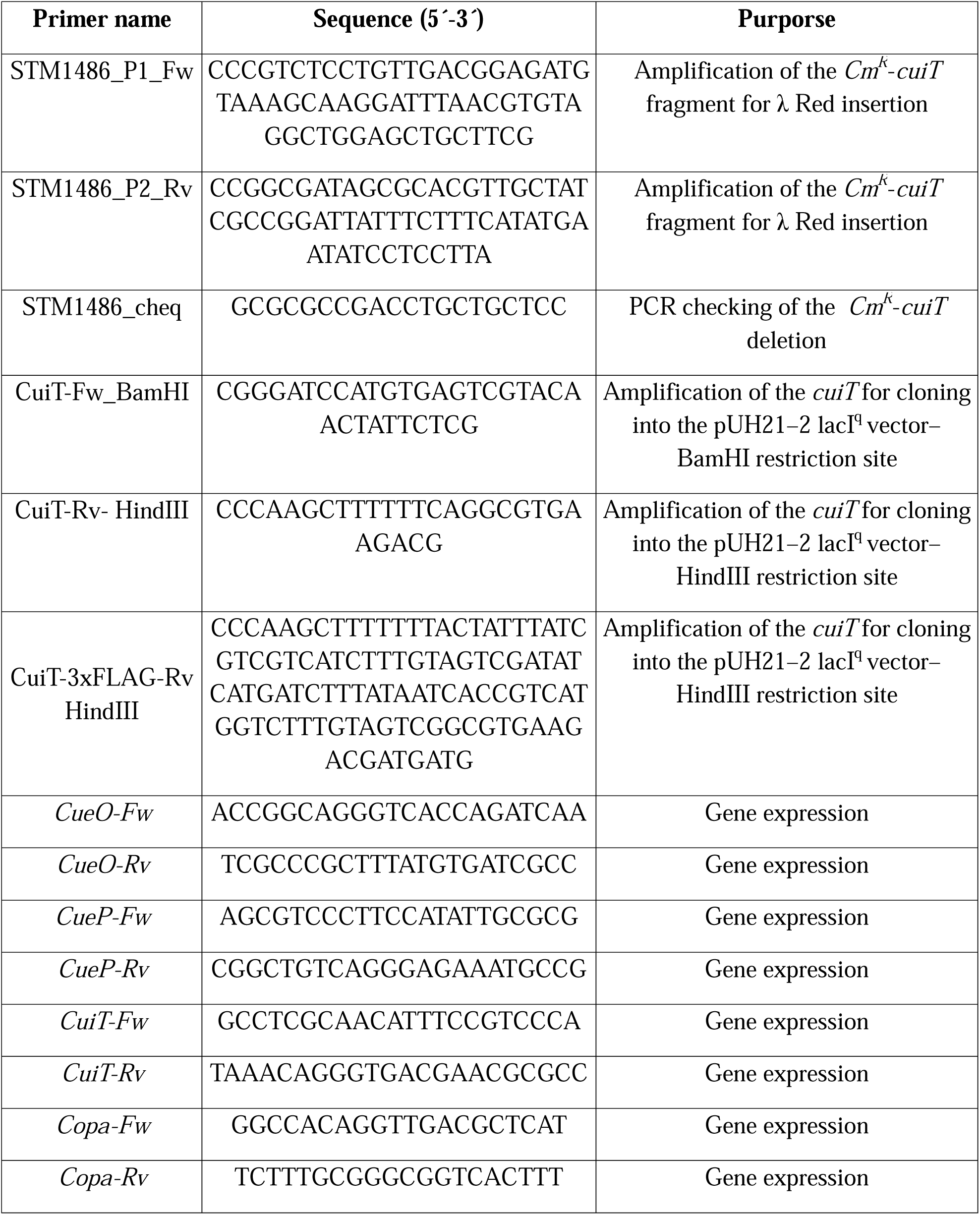
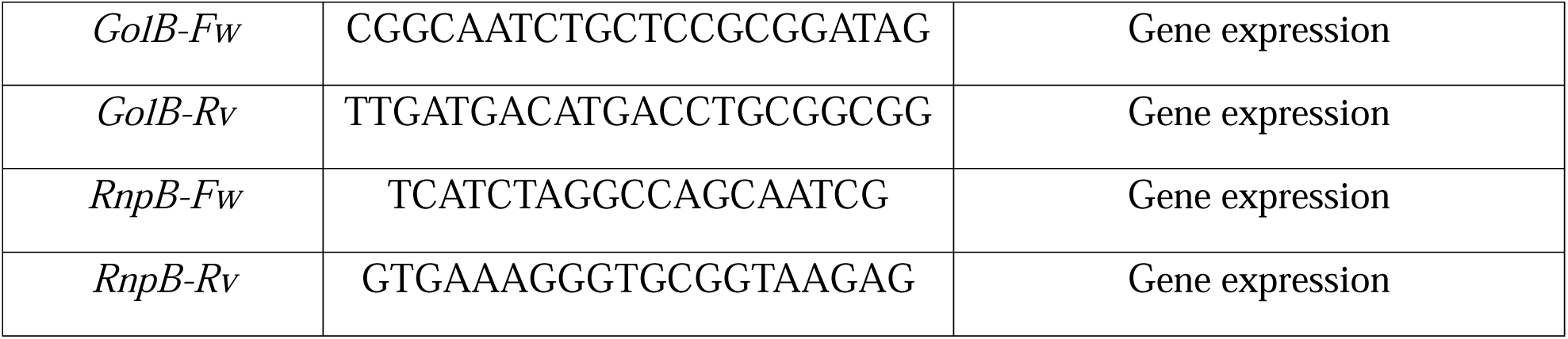
Oligonucleotides used in this study.

